# Impact of flagellated and elongated morphological phenotypes on the focusing behaviours of biological cells in inertial microfluidic devices

**DOI:** 10.1101/2024.05.16.594321

**Authors:** Jessie Howell, Nicole Hall, Sulochana Omwenga, Tansy C. Hammarton, Melanie Jimenez

## Abstract

Inertial microfluidics has demonstrated tremendous potential to impact biological - and notably medical - fields, by offering a highly versatile, portable and cost-effective approach to cell focusing and sorting. While the range of applications of inertial devices spans medical diagnostics, bioprocessing or water engineering to mention a few, translation is still impeded by the lack of clear understanding of cell interactions in such devices. This often leads to bespoke designs that take years of development and characterisation for one targeted application, and limited tools for informed optimisation. A more fundamental knowledge of inertial behaviours is key to future translational works and impact, by enabling a deeper understanding of inertial forces in biological systems. Towards this goal, this paper focuses on high-throughput morphological phenotyping of the single-celled, flagellated parasite *Leishmania mexicana* to better understand how variations in cell body length, width and flagellated status impact the focusing patterns of highly non-spherical cells in curved inertial devices. Some of the key findings in this study include i) not all organelles, such as flagella, will alter focusing if the body shape is conserved, ii) the impact of cell shape is specific to a channel design and slight changes in e.g., cell confinement can completely change focusing patterns, iii) elongated prolate-like cells align in different orientations depending on their lateral position with a curved channel and iv) despite variabilities observed in focusing patterns for elongated versus rounder cell phenotypes, large morphological variations can be completely overcome at high Reynolds numbers so that all phenotypes tightly focus at a single position (here towards the channel outer wall). This last finding, in particular, may open new avenues for highly efficient cell enrichment processes, such as for the detection of pathogens in water.

## Introduction

Inertial microfluidics is an attractive method for sorting particles, offering a high-throughput, label-free and passive method of separation. While it has been commonly described for sorting cells based on their morphological and mechanical phenotypes, particle size – and more specifically their equivalent diameter - remains the most widely considered factor for driving inertial microfluidic separation and related empirical correlations [1-3]. Biological cells with a round morphological phenotype are often hypothesised to experience inertial forces in a similar fashion to size-matched rigid spherical (*e*.*g*., polystyrene) beads, which are consequently used as model systems for design development [4]. Deformability has also been shown to strongly affect focusing behaviours, where the presence of a deformability-induced lift force causes the migration of deformable objects away from more rigid particles [5-7]. This sorting parameter has been successfully applied to *e*.*g*., the separation of deformable cancer cells from blood, diagnosis of malaria through identification of infected red blood cells (RBCs) or the purification of manufactured RBCs [7-9]. In contrast, the contribution of shape is less well documented.

Initial work on spheroid particles in straight inertial microfluidic (IM) channels (*i*.*e*. without the presence of Dean forces) has reported complex rotational behaviours, with prolate (elongated ellipsoid) particles and oblate (disc-like ellipsoid) particles showing rotations described as “kayaking”, “log-rolling” and “tumbling” depending on their position in the flow and the strength of the inertial forces. When these particles focus within the flow, they tend to exhibit a stable rotation with prolate particles “tumbling” and oblate particles “log-rolling” [10-12]. By increasing the Reynolds number further, particles have been shown to align with the flow and rotate periodically or stop rotating altogether, although this behaviour has not been observed in all studies [11, 13]. The longest dimension of a particle (rotational diameter) has been reported as a measurement equivalent to the diameter of a spherical particle, with non-spherical and spherical particles showing a high correlation independent of their shape [10, 14, 15]. Importantly however, this pattern does not translate to all non-spherical shapes, *e*.*g*. non-symmetrical ones, as described elsewhere [16]. It can also be noted that most studies to date have investigated the role of shape in straight IM channels, with fewer systematic explorations in curved designs. Although the relevance of the equivalent diameter has also been shown experimentally for non-spherical objects in Dean flows, divergence in focusing behaviours for differently shaped (but volume- and dimension-matched) beads highlights a higher complexity in fluid-particle interactions compared to straight IM devices [15]. These differences have been partially linked to particle rotation but also to the generation of secondary vortices which can push non-spherical objects away from traditional focusing positions (*e*.*g*., away from the inner wall in standard spiral designs) [17]. Since shape matters in biology [18], IM devices have been applied to a range of non-spherical cellular systems. Using a spiral IM channel with a trapezoidal cross section, small and round *Caenorhabditis elegans* eggs could be isolated from later development stages with a vermiform shape [19]. Keinan showed that sorting of budding yeast in serpentine channels could be used to identify different populations based on cell age [20]. Yeast has also been separated based on shape into different cell cycle stages, obtaining up to 94% purity for singlet cells and 31% purity for budding stages [14]. Beyond yeast, a range of designs have been engineered to sort differently-shaped microalgae [21-23] and spermatozoids [24-26], to mention a few.

Importantly, cells typically present biological variations in their shape; replicative cells, for instance, undergo a continuum of alterations to their size and shape throughout their cell cycle. An accurate representation of the effect of cell shape in IM devices can consequently be challenging to obtain, making robust conclusions on focusing patterns harder to draw. In this work, we investigate more closely the impact of the morphological phenotype on focusing patterns in curved IM devices. More specifically, high-throughput imaging flow cytometry was used to characterise populations of the naturally elongated prolate-like parasite *Leishmania mexicana* that had been differentiated or chemically treated to vary in size, shape and/or flagellated status. Single cell high-speed imaging in curved IM devices was then deployed to demonstrate that although shape plays a critical role in focusing patterns, large differences in size and/or shape can be completely circumvented for precise focusing to a single lateral position within the channel. In contrast to prior studies, it is also shown here that the presence of a flagellum has little impact on focusing in the tested conditions [25]. These findings pave the way to a better understanding of focusing in IM devices for non-spherical cells.

## Materials and methods

### Cell preparation

*Leishmania mexicana* M379 Cas9 T7 promastigotes [27] were cultured according to [28] in medium 199 (Gibco™) supplemented with 10% foetal bovine serum (FBS; Gibco™), 26.2 mM sodium bicarbonate, 0.005% haemin, 40 mM 4-(2-hydroxyethyl)piperazine-1-ethanesulfonic acid pH 7.4, 32 µM hygromycin and 50 µM nourseothricin sulphate at 27°C. Analysis was carried out on cells in their logarithmic stage of growth (1 x 10^6^ - 7 x 10^6^ cells.ml^-1^). Cells were deflagellated using a modification of the protocol outlined in [29]. Cells were fixed in 4% paraformaldehyde (PFA) in phosphate buffered saline (PBS) for 15 minutes at 4°C and washed 3X in PBS, before being resuspended in 5 ml of a solution of 10 mM 1,4-piperazinediethanesulfonic acid (PIPES), 1 mM CaCl_2_, 1 mM MgCl_2_ and 0.32 M sucrose adjusted to pH 7.2. CaCl_2_ was added to a final concentration of 0.075 M, and the sample was kept on ice. Flagella were removed from the cell bodies by drawing the resuspended cells through a 10 ml syringe fitted with a gel loading pipette tip 200 times. Sucrose gradient centrifugation was carried out as described in [29] to isolate the deflagellated cell bodies from cells still possessing a flagellum, cell fragments and sheared flagella. The cell body fraction was washed with PBS prior to use in imaging flow cytometry or inertial microfluidic experiments. Axenic amastigotes were generated by culturing mid-log phase M379 Cas9 T7 cells in Schneider’s Drosophila medium (Gibco™) pH 5.5, supplemented with 10% FBS at 32°C for at least 72 hours. To synchronise cells in different cell cycle stages, mid-log phase parasites were incubated with either 5 mM hydroxyurea (Sigma) or 5 µM flavopiridol (Selleck Chemicals) for 10 hours before being harvested and washed in PBS. Cells were then fixed with 4% PFA/PBS for 15 minutes at 4°C and washed 3X in PBS prior to testing.

### Cell characterisation using imaging flow cytometry

Morphological analysis of both live and fixed cells was carried out using imaging flow cytometry. Samples were resuspended in 60 μl PBS prior to acquisition on a Cytek® Amnis® ImageStream®X Mk II. ∼30,000 events were acquired for each sample using INSPIRE^™^ software, collecting data from the brightfield channel using the 60X objective lens. The subsequent files were analysed using IDEAS® software to extract morphological measurements of individual cells. The cell body mask and the gating strategy outlined in [30] were used to remove images containing speed beads (added to the sample during acquisition for calibration purposes), debris, cell rosettes, two cells and out of focus cells. Area, aspect ratio (the minor axis of a fitted ellipsoid *vs* the major axis), length and width measurements were calculated for the remaining population of cells and the data was exported from IDEAS® for further analysis in R. Data smoothing was carried out for the length and width measurements to account for the relatively low resolution of the images (length and width measurements are calculated in IDEAS® in increments of 0.33 μm and 0.66 μm, respectively). For this, a randomised number between -0.33 and 0.33 was added to each length measurement, and between -0.66 and 0.66 for width measurement.

### Bead specifications

Various shapes and sizes of polystyrene beads were tested throughout this work. Spherical beads (3 and 5 μm diameter) were sourced from Magsphere Inc. along with 2.6 x 3.2 μm^2^, 2.5 x 4.0 μm^2^, and 3.8 x 5.1 μm^2^ pear-shaped beads and 2.8 x 4.0 μm^2^, 4.5 x 6.3 μm^2^, 5.1 x 7.7 μm^2^ peanut-shaped beads. Spherical beads (4, 6, 8 and 10 μm diameter) were sourced from Invitrogen.

### Microfluidic setup

The microfluidic chips (Epigem) used in this work consisted of a 6-turn Archimedean spiral with an inlet and four outlets. Two sizes of chips were used with a channel height and width of either 30 x 170 µm^2^ or 60 x 360 µm^2^. Both bead and cell samples were prepared in PBS to a concentration of 5 x 10^5^ particles.ml^-1^ for IM analysis and injected into the smaller device at flow rates of 0.2 – 1.5 ml.min^-1^ (Re = 33.3 – 250) or the larger device at 0.3 – 4.0 ml.min^-1^ (Re = 23.8 – 317.5). A Nemesys high pressure pump (Cetoni) was used to pump the samples through 1/16′′ polytetrafluoroethylene (PTFE) tubing with an internal diameter of 0.5 mm. Particle focusing behaviour was analysed using an AcCellerator cell analyser (Zellmechanik), which consisted of a high-speed camera and an inverted microscope. The chips were imaged using either a 20X or 10X objective for the smaller or larger device respectively. The focusing position of the particles was acquired using the AcCellerator cell analyser’s Shape-In (version 2.5.5.815) software to acquire images of each cell within the channel at 100 fps. Analysis was carried out in Shape-Out 2 (version 2.13.6) to extract the position of the particles within the channel, which were subsequently analysed and plotted in R.

### Deformability measurements

Deformability measurements were carried out using an AcCellerator cell analyser as outlined in [31]. Sheath fluid (CellCarrierB, Zellmechanik) and PFA-fixed flavopiridol-treated cells resuspended in CellCarrierB were injected into a 30 x 30 μm^2^ microfluidic chip at flow rates of 0.24 μl.s^-1^ and 0.08 μl.s^-1^, respectively. 1000 events were acquired using a frame rate of 300 fps using Shape-In. Measurements were taken in triplicate in both the measurement channel and the channel reservoir (control). Analysis was performed in Shape-Out 2 firstly to remove debris (e.g. fragments of cells), clumps of cells, and events with incorrect contouring. This was achieved by gating the events based on the porosity of the object contour *vs* its area. The subsequent gate was applied to both the measurement and the channel reservoir data, and the resulting populations were plotted using the area of each object *vs* its degree of deformation. The density of the data points was calculated using a Gaussian kernel density estimate, which was used to generate contour plots showing the 95^th^ and 50^th^ percentiles of the data.

## Results

### Impact of flagella on focusing patterns

As various studies have looked at IM devices for sorting cells based on their cell cycle stage, promastigote *Leishmania mexicana* M379 Cas9 T7 parasites were used as the biological model in this study. When cultured *in vitro*, this parasite typically forms mixed populations of cells which vary in size and shape according to their stage in the cell cycle (G1, S, G2, mitosis (M) and cytokinesis) [30, 32]. This highly non-homogeneous population is named hereafter the parental cell population and was used as the control for all tests carried out in this study. Since this parasite is naturally flagellated, we first investigated whether the presence of a flagellum can alter focusing patterns. Two approaches were considered for this – chemical deflagellation and differentiation. Chemical deflagellation was carried out in line with previous work [29] while differentiation was used to generate axenic amastigotes. Amastigotes are the mammalian-infective stage of the parasite, which naturally possess only a vestigial flagellum presenting as a small bulbous tip that emerges from the cell body, yet the internal structures associated with this organelle are maintained [33]. Since these cells are highly variable in their shape, the use of imaging flow cytometry (as opposed to *e*.*g*., standard microscopy) was essential to characterise their profile in a meaningful manner. **Figure 1a** depicts examples of brightfield images collected with the imaging flow cytometer for the three populations, with the flagella clearly visible for the parental promastigotes. Analysis of thousands of cell images confirmed that chemically deflagellated cells had a similar range of morphologies to parental cells (**Figure 1c**), with a maximum deviation of 0.12 from 0.5 when comparing the area under the curve (AUC; **Figure 1b**). In contrast, as reported in the literature [34], amastigotes were significantly shorter (average length of 5.5 μm, compared to 9.5 μm for parental) and rounder (mean aspect ratio of 0.63, compared to 0.35 and 0.41 for parental and deflagellated parasites, respectively) with a width comparable to that of parental cells (average width of 3.5 µm, compared to 3.3 µm for parental). This is similarly demonstrated by the AUC values, in which area, aspect ratio and length demonstrate high deviations from 0.5.

**Figure 1.**
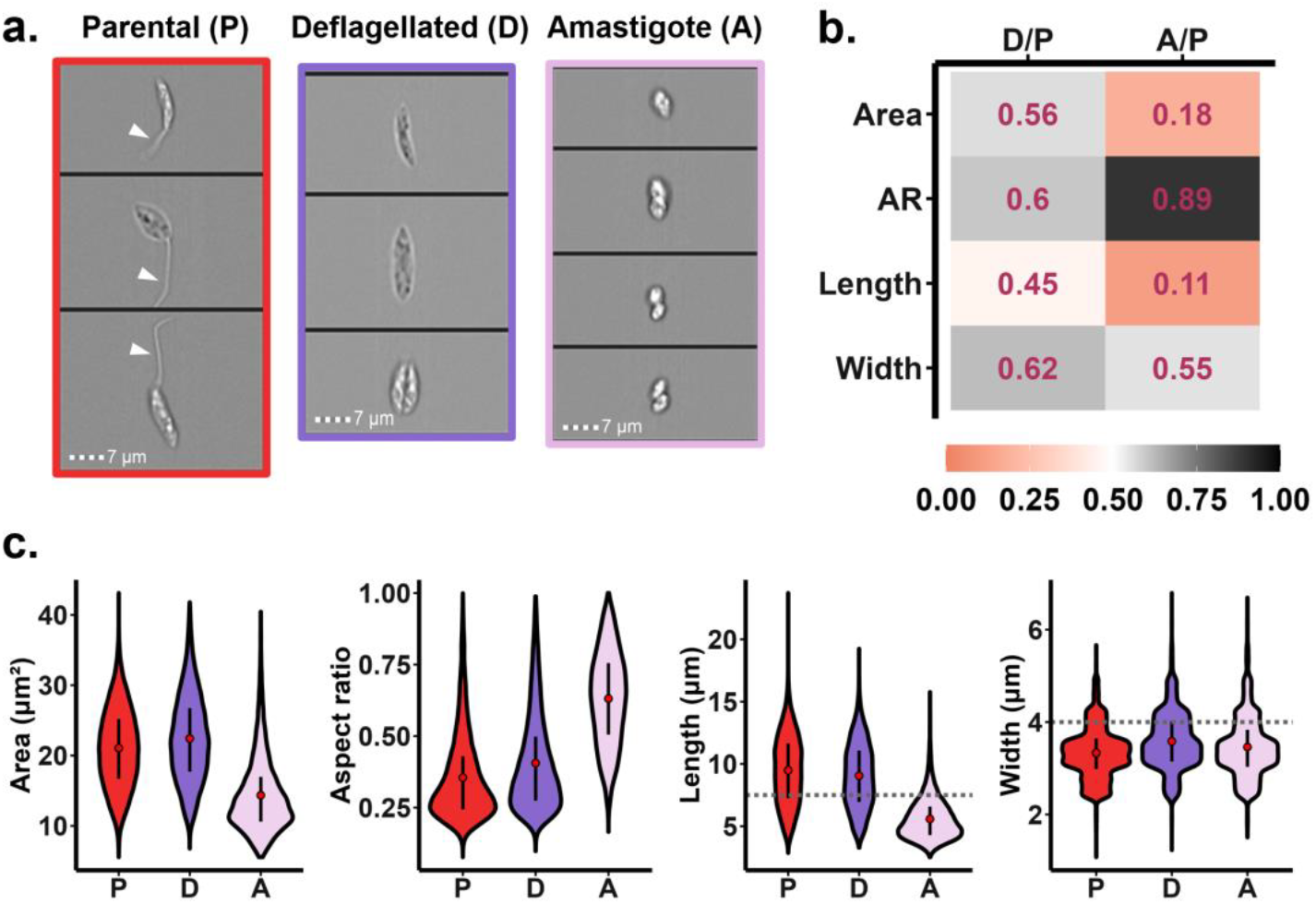
Characterisation of the morphological phenotype of fixed parasites with and without flagella using imaging flow cytometry. (a.) Example images of fixed parental (P), deflagellated (D) and amastigote (A) cells. The white arrowheads identify cell flagella. Scale bars: 7 μm. (b.) Comparison of the area, aspect ratio (AR), length and width distributions between deflagellated and parental samples (D/P), and amastigote and parental cell population (A/P), as calculated from the area under the curve. A score close to 0.5 represents a distribution with a high similarity to the parental cell population, while a score closer to 0 or 1 denotes a high deviation in the distribution of measurements. A value < 0.5 or > 0.5 indicates that the morphological measurements are smaller or larger than the parental cells, respectively. (c.) Violin plots depict the area, aspect ratio, length, and width distributions of the different cell populations. The red circles show the mean value, the black bars the 25^th^ to 75^th^ percentiles and the dotted grey lines indicate the thresholds applied to the length and width measurements (7.5 μm and 4.0 μm, respectively; *n* ≥ 12,962).

While considerable variations in size and shape were observed within all cell populations (for example, the length of parental cells ranged from ∼ 3 to 15 µm, reflecting their cell cycle stage heterogeneity), the majority (∼60.7%, **Figure S1**) of parental and deflagellated cells were long (> 7.5 μm in length) and narrow (< 4 μm in width), while amastigotes were predominantly (73.5%) short (≤ 7.5 μm in length) and narrow. These thresholds were chosen to be relevant to the dimensions of the 30 x 170 μm^2^ channel used: 7.5 μm corresponds to ¼ of the channel height (∼ half of a Dean vortex, assuming two vortices) while a width of 4 μm provides a particle confinement ratio (λ) > 0.07 [35].

To characterise the focusing behaviour of these three cell populations, single cell high-speed imaging was performed near the outlet of a spiral IM channel. In such a design (**Figure 2a**), spherical rigid particles with an equivalent diameter of 10 μm typically focused at the inner wall for the range of Reynolds numbers investigated in this work (up to Re = 250.0, **Figure S2**) and data are consequently presented as distances from the inner wall. While some enrichment was seen at the inner wall for the parental population at Reynolds numbers of 83.3 and 116.7 (**Figures 2b** and **c**), both the inherently elongated parental and deflagellated parasites predominantly focused close to the outer wall for Reynolds number of 116.7 and above, which confirmed that in a curved channel, a particle’s rotational diameter is not equivalent to the diameter of a spherical particle as has been previously reported [14, 35]. Instead, this indicated an increased contribution of shape on focusing patterns. Furthermore, few differences were observed in the focusing behaviour between the parental and deflagellated parasites, indicating that the presence of a flagellum, for a conserved morphology, does not significantly alter focusing patterns for the tested conditions.

**Figure 2.**
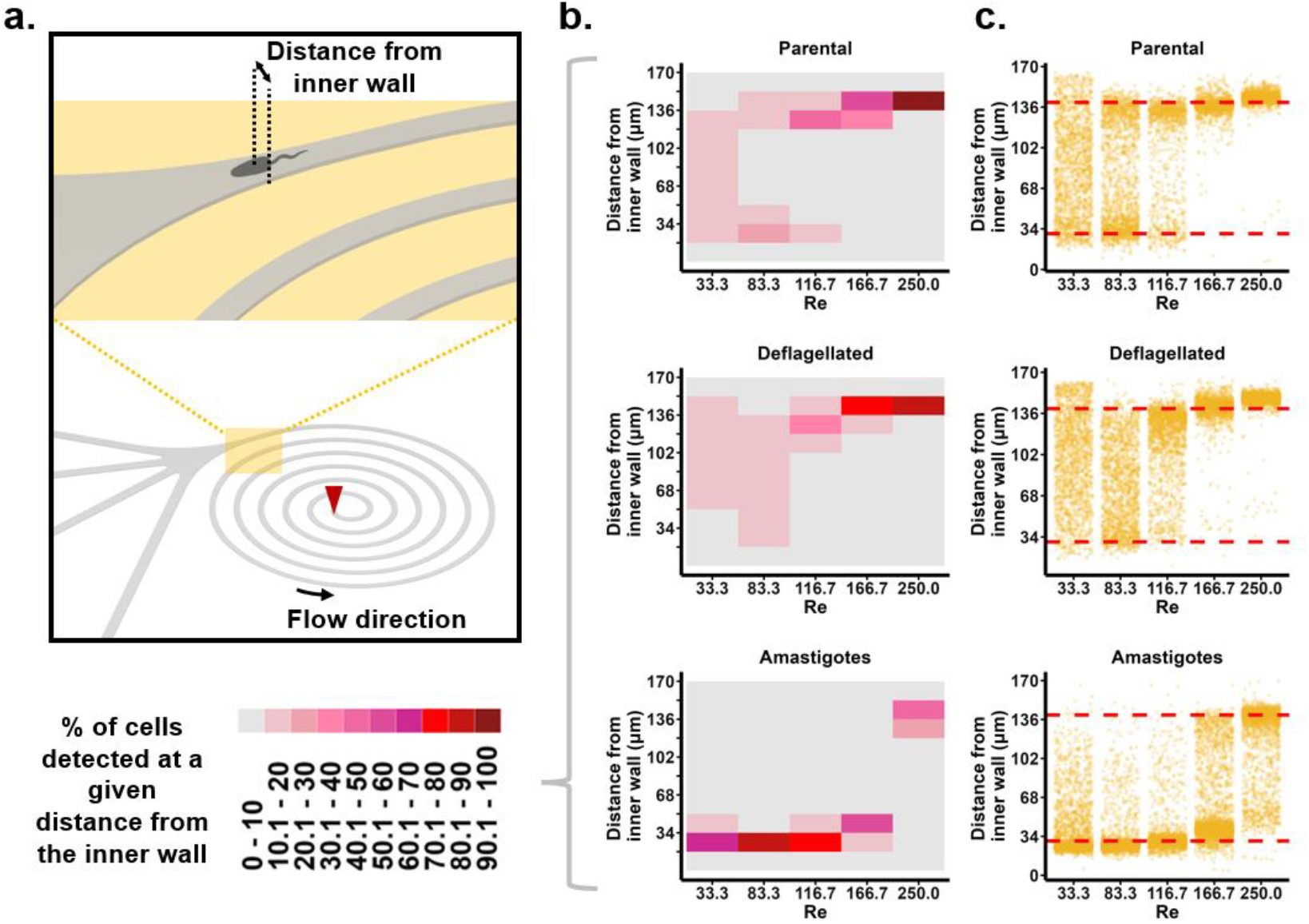
Characterisation of the focusing behaviour of fixed parental, deflagellated and amastigote cells in a spiral inertial device. (a.) Schematic of the inertial spiral channels used in this study (either with a cross section of 30 x 170 µm^2^ or 60 x 360 µm^2^), with a zoomed-in view near the outlet. The red arrow depicts the sample inlet and fixed parasites were imaged next to the opening for the outlet. Single cells were imaged using a high-speed camera and their location within the image was automatically extracted using Shape-In software. Their relative position within the channel was thus calculated as the distance from the inner wall (max distance = 170 µm or 360 µm). (b.) Percentage of parasites (*n* ≥ 965) focusing at a given distance from the inner wall of the 30 x 170 μm^2^ device (channel width has been divided into 10 sections), with red zones highlighting a high focusing concentration with >70% of cells found within a tenth of the channel width. (c.) Representation of focusing patterns with individual dots corresponding to a single parasite (*n* ≥ 965) detected in the region of interest. The red dashed lines indicate a distance from the inner wall of 30 and 140 µm as a reference. Re denotes the Reynolds number.

In contrast, amastigotes with a short, narrow and higher aspect ratio morphology (a morphology closer to that of spherical beads) focused tightly to the inner wall at lower Reynolds numbers (≤166), with >80% of cells focusing within a 17 μm band at Re = 83.3 (**Figure 2b**). As the Reynolds number increased, the cells were seen to shift towards the outer wall. Interestingly, for the highest tested Reynolds number (Re = 250), ≥60% of cells from each population analysed (parental, deflagellated and amastigote) were concentrated within a 34 µm (119 – 153 μm from the inner wall) region close to the outer wall - irrespective of their significant morphological differences.

To further validate the effects of shape on sorting, live parental cells were sorted at Re = 116.7 into four outlets and the morphology of the cells in each outlet was analysed (**Figure S3a**). From these data, the biggest change in the distribution of morphologies, when compared to the unsorted control, was seen at the inner wall for aspect ratio (AUC of 0.63; **Figure S3b**). This corresponded to an enrichment of short and wide cells (i.e. a higher aspect ratio) at the inner wall (17.1% at the inner wall *vs* 9.2% in the unsorted control; **Figure S3c**). Similarly, analysis of the percentage of cells sorted into each outlet confirmed that short and wide cells were enriched at the inner outlet (≥12.4% increase compared with any other outlet) while short and narrow cells were evenly distributed between the outlets at the inner and outer wall (30.4% vs 27.0%, respectively; **Figure S3d**). Long and narrow cells on the other hand were predominantly found at the outlet closest to the outer wall (45.1%), with only 15.6% at the inner wall. These data are not only consistent with previous findings on the impact of shape on focusing patterns but may also indicate intricate cell-cell interactions that can impact sorting as previously demonstrated for mixed cell populations [36].

### Impact of cell shape on focusing patterns

We then explored whether the shared focusing position at high Reynolds number observed for the three previous cell populations was translatable to a wider range of cell shapes. Towards that goal, two additional cell populations were tested, with their characterisations presented in **Figure 3**. Parental parasites were chemically treated to arrest cells at specific stages (and consequently morphologies) of the cell cycle. Flavopiridol-treated cell populations – mostly arrested in G2 phase [30] – had a similar distribution of length to the untreated parental cells (mean length of 10.0 µm, compared to 9.5 µm for parental with an AUC of 0.54) but were generally wider (mean width of 4.9 µm, compared to 3.3 µm for parental and an AUC of 0.92). Hydroxyurea-treated cell populations on the other hand, reported to arrest other *Leishmania spp*. at G1/S [37, 38], had a highly similar distribution of aspect ratio sizes, with an AUC of 0.48 compared to the parental, but had an increase in the proportion of longer and wider cells (**Figure 3b and S1**).

**Figure 3.**
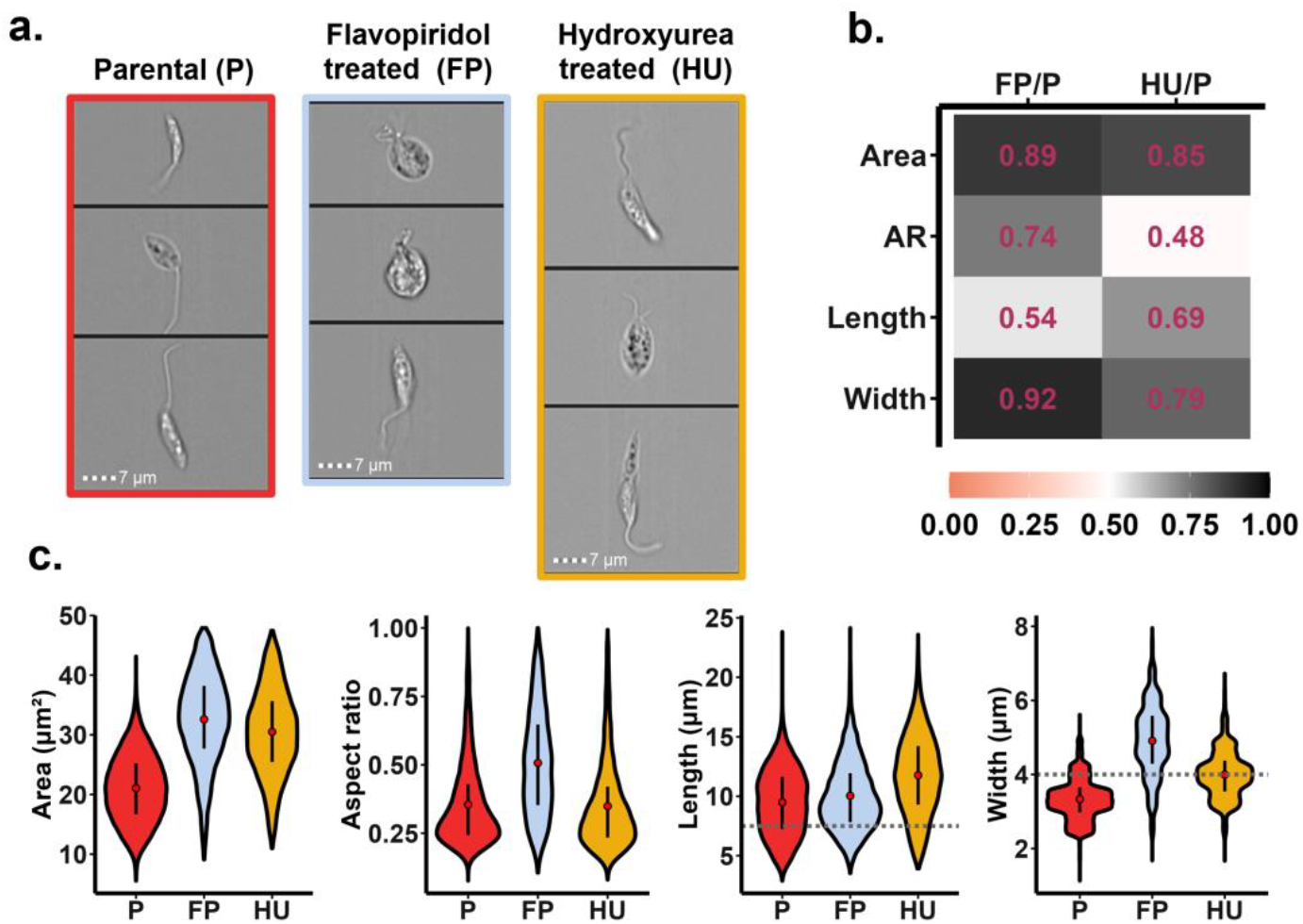
Characterisation of the morphology of cell cycle-arrested *L. mexicana* using imaging flow cytometry. (a.) Example images of fixed parental (P), flavopiridol-treated (FP) and hydroxyurea-treated (HU) cell populations. Scale bars: 7 μm. (b.) Comparison of the area, aspect ratio (AR), length and width distributions between FP-treated and parental cells (FP/P), and HU-treated and parental (HU/P), calculated as the area under the curve. A score close to 0.5 represents a distribution with a high similarity to the parental cell population, while a score closer to 0 or 1 denotes a high deviation in the distribution of measurements. A value < 0.5 or > 0.5 indicates that the measurements are smaller or larger than the parental cells, respectively. (c.) Violin plots depict the area, aspect ratio, length and width distributions of the different cell populations with the red circle showing the mean value and the black bars the 25^th^ - 75^th^ percentiles (*n* ≥ 12,962).

Looking at the focusing position of parasites with these modified cell morphologies, different trends emerge (**Figure 4**). To analyse cells with a higher rotational diameter but with a similar aspect ratio, hydroxyurea-treated cells were compared to the parental population. At Re = 83.3 and 116.7, both populations concentrated at two concurrent focusing points, namely ∼30 µm from the inner and outer wall (**Figure S4**). Interestingly, a higher proportion of hydroxyurea-treated cells focused towards the inner wall at these same flow rates. This was unexpected as in **Figure S3c** it was seen that longer parental cells predominantly focused to the outer wall. The increase in the proportion of long hydroxyurea-treated cells at the inner wall may be explained by differences in particle rotation: when comparing images taken with the high-speed camera, elongated cells that focused towards the inner wall predominantly adopted an orientation perpendicular to the channel walls, while cells focusing towards the outer wall seemed to align with the flow (**Figure S5**). It is thus hypothesised that a low aspect ratio is required for particles to both align with the flow and focus to the outer wall, and above a certain confinement, particle rotation is modified such that particles maintain a horizontal alignment and focus to the inner wall. Elongated cells therefore may exhibit different focusing patterns, either at the inner wall or outer wall, depending on the Reynolds number, their aspect ratio, and their confinement within the channel.

**Figure 4.**
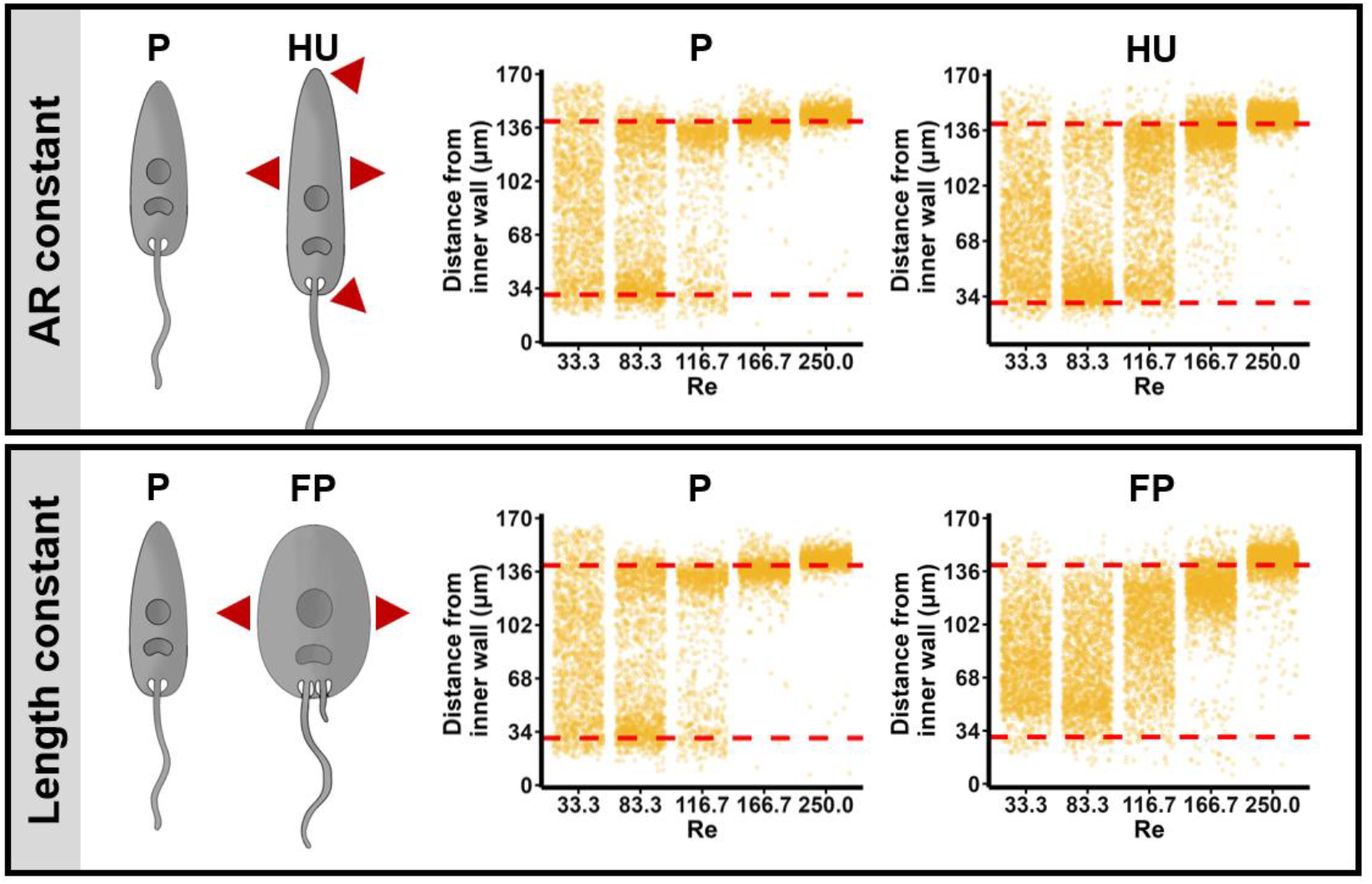
Characterisation of the focusing behaviour of fixed parental, flavopiridol-treated (FP) and hydroxyurea-treated (HU) cell populations in a spiral inertial device. HU-treated cells (top panel) and FP-treated cells (bottom panel) are compared to parental cell populations. A graphical representation is given on the left of each panel, with the red arrows indicating the dimensions that were modified for both cell populations in comparison to the parental parasites. Single cells (n ≥ 965) were imaged in the 30 x 170 μm^2^ device using a high-speed camera and their position near the outlet were quantified as a distance from the inner wall (max distance = 170 µm). The red dashed reference lines indicate a distance of 30 µm from both the inner wall and outer walls. Re denotes the Reynolds number and the same graph for P is displayed for each panel.

In contrast, increasing the width but not the length of cells (thereby increasing the aspect ratio) seemed to result in the parasites remaining dispersed throughout the channel instead of migrating to either wall, as seen for flavopiridol-treated cells at Re ≤ 116.7 (**Figure 4 and Figure S4**). This contradicts the literature which has reported that the length and not the width of non-spherical beads has an effect on focusing [14], further suggesting that aspect ratio is an important consideration for the focusing of non-spherical cells. Similarly to the other cell types tested, at high Reynolds numbers, both the hydroxyurea- and flavopiridol-treated cells occupied the same focusing position within ∼30 μm from the outer wall.

We finally wanted to compare these trends to rigid model particles as done in other studies [20, 22, 39]. We aimed to match these objects as closely as possible to some of the biological cells tested in this study, although the range of shapes available was manufacturer dependent. A summary of the different morphologies tested is given in **Figure 5**, with bead focusing behaviours detailed in **Figure S6**. As shown in these figures, all sizes of the rigid objects tested (between 3 to 10 µm in diameter) focused tightly to the inner wall and none emulated the behaviours observed with parasites at the outer wall. Even closely matched particles (*e*.*g*., amastigotes with pear-shaped beads) diverged in focusing behaviours, especially at the highest Reynolds number tested where virtually no focusing within 30 μm of the outer wall was observed for any beads. Peanut-shaped beads (aspect ratio ∼0.70) showed the furthest migration to the outer wall, but the focus was significantly more spread than the one observed for parasites (with or without flagella). As described in other studies, closely matched beads with different shapes (such as those highlighted by blue arrows in **Figure 5**) were also seen here to behave differently from each other [15]. These data confirm that shape plays a critical role in focusing patterns.

**Figure 5.**
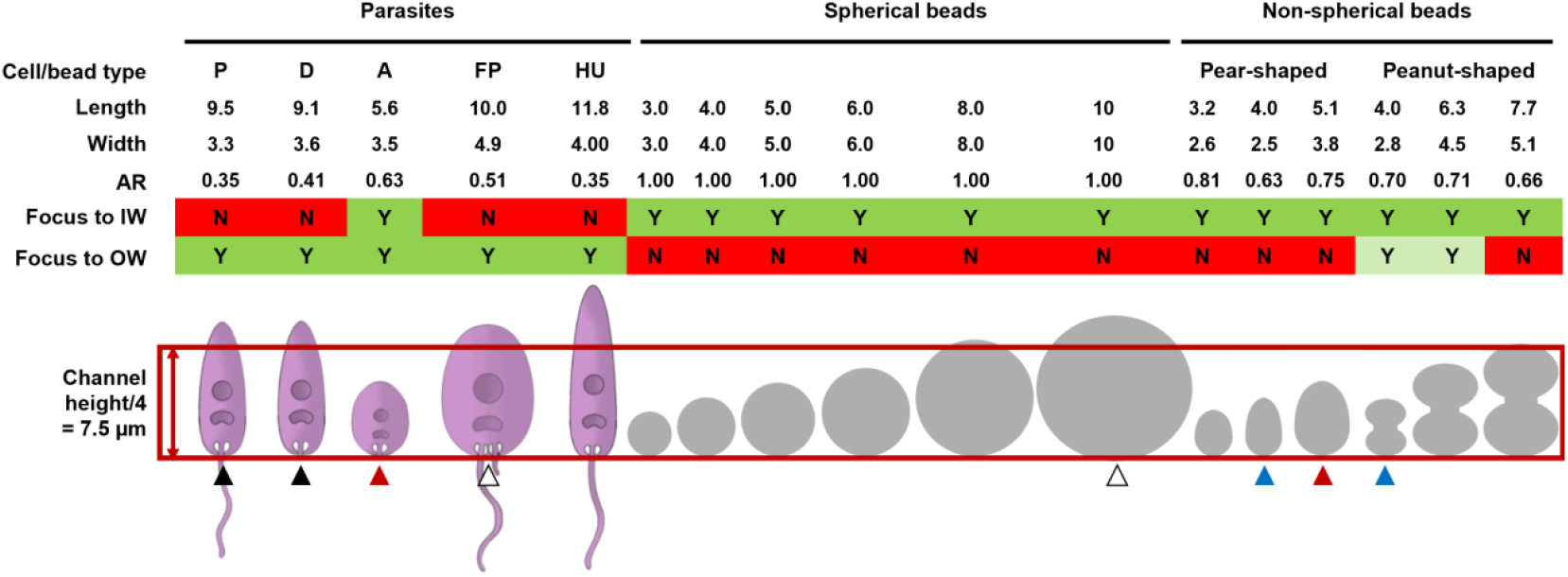
Summary of morphological phenotypes tested in this work. Objects are characterised in this schematic by their average length (longest dimension, µm), width (shortest dimension, µm), and aspect ratio (AR). P, D, A, FP and HU denote parental, deflagellated, amastigote, flavopiridol- and hydroxyurea-treated cells, respectively. A green Y (Yes) or red N (No) indicates whether particle focusing was observed for at least one flow rate at 30 μm from the inner wall (IW) or 30 μm from the outer wall (OW). A light green Y denotes a focusing position within the outer half of the channel, but further than 30 μm from the outer wall. The morphological phenotype of particles was also compared to the channel quarter height (corresponding to half a Dean vortex). Arrows depict objects with the most similar morphologies: black for parental and deflagellated cells, red for amastigotes and pear-shaped beads, white for FP-treated and a spherical bead, and blue for pear- and peanut-shaped beads.

## Discussion

This study aimed to explore the role of cell shape in inertial focusing. Although size and shape have both been exploited as sorting parameters in IM devices, one of the key challenges to better interpret results is the intrinsic variability in morphologies encountered in most biological systems. Without high-throughput characterisation of the morphological phenotype for a given cell population, conclusions on inertial focusing behaviours can be skewed or biased. Here we used high-speed imaging to not only characterise single cell behaviours in IM devices, but also to quantify morphological variations with high accuracy.

Initial tests with the parental cell population demonstrated that the elongated prolate-like *L. mexicana* displayed a focusing position towards the outer wall at Reynolds numbers as low as 83.3. In the literature, a preferential focusing at the outer wall has been observed in such devices for deformable cells such as cancer or red blood cells [5, 9]. Here, cell deformability was not expected to be the primary reason for this migration to the outer wall; both live and fixed cells exhibited the same focusing behaviour despite fixation being known to reduce cell deformability. However, it can be expected that the fixed cells were still not as rigid as *e*.*g*., polystyrene beads, thus deformability cytometry [31] was attempted to quantify the mechanical properties of cells. This proved challenging with the elongated shapes studied but data for rounder flavopiridol-treated cells nonetheless confirmed that cells could alter their shape under controlled shear forces even with fixation (**Figure S7**). It is consequently important to acknowledge that deformability may play a role in the findings of this work. It is also possible that the migration of both elongated cells and deformable particles towards the outer wall at high flow rates occurs under the same mechanism. The mechanisms driving the migration of deformable particles to the outer wall are still not understood; however, it is known that deformable particles experience a shape change at high shear rates to take on an elongated morphology [16]. Furthermore, instead of rotating with the flow, these particles have been described to tank tread with the particle thus maintaining a consistent deformed shape and behaving similarly to non-rotating elongated parasites.

Beyond deformability, focusing at the outer wall has also been attributed in the literature to the presence of flagella. Flagellated human spermatozoids were observed to shift towards the outer wall, while deformable red blood cells or deflagellated spermatozoids remained at the inner wall [25]. It was consequently concluded that the presence of flagella could alter the rotation of particles and induce migration. In contrast, here we demonstrated that removal of the flagella had minimal impact on the focusing of *L. mexicana* parasites. There are two possible explanations for the discrepancy in results. The preferred hypothesis, consistent with the work on spermatozoids, is that a lack of particle rotation causes the migration to the outer wall. For the spermatozoid cells, this is induced by the presence of flagella, and on removal of the flagella the resulting cell bodies (which are closer to spherical) are capable of rotating in the flow and thus focus to the inner wall. In contrast, for the tested *L. mexicana* cells it may be that both the elongated shape of the cells and the flagella are analogous in preventing particle rotation and thus by removing the flagellum, the elongated shape still prevents particle rotation. The second hypothesis is that treatments such as sonication (used to deflagellate spermatozoids in [25]) can significantly alter the morphology of cells making isolating the impact of flagella challenging [40]. Using imaging flow cytometry, we employed a protocol for deflagellating our parental parasites without drastically changing the dimensions of the cell body. In the tested conditions, we could not induce focusing at the inner wall for deflagellated cells; only when the cell body morphology was altered, such as with smaller and rounder amastigote cells, did focusing at the inner wall occur at lower Reynolds numbers.

Previous research carried out with heterogenous microalgae in straight IM channels showed that elongated cells remained close to the channel centre while rounder cells drifted closer to the outer walls [23]. Assuming these patterns are conserved in a curved IM device, and assuming two counter rotating Dean flows within the channel’s cross section [41], elongated cells locating closer to the centre line would be dragged by Dean forces toward the outer wall as observed for deformable objects [5, 9]. Sorting of parental cells confirms this trend in the spiral IM device, showing an increased concentration of rounder cells closer to the inner wall and of narrower/longer cells closer to the outer wall (**Figures S3c and d**). Importantly however, and similarly to findings regarding cell length, a rounder morphology is not the sole condition for focusing towards the inner wall as rounder flavopiridol-treated cells located towards the channel centre. Instead, specific confinement may be required for focusing at the inner wall. In a channel with a larger cross section, parental parasites are indeed seen to focus towards the inner wall at lower Reynolds numbers, while maintaining a focusing position at the outer wall at higher flow rates (**Figure S8**). Aspect ratio and confinement consequently both play an important role in the focusing of non-spherical cells at both the inner and outer wall. It is expected that these morphological features affect particle rotation within the curved channel which in turn leads to altered focusing positions, as was observed between parental and hydroxyurea treated cells[11].

While most studies with IM devices aim to sort populations with different morphologies, an interesting finding of this work is the ability to concentrate a widely heterogeneous population into a single, narrow focusing stream at the outer wall. All the *L. mexicana* populations tested in this work - irrespective of their flagellated status, shape or chemical treatment – were observed to focus at the same equilibrium position ∼ 30 µm away from the outer wall (distance equivalent to the channel height) at a high Reynolds number (∼250.0). These findings were replicated in a larger 60 x 360 µm^2^ channel, with cells focusing ∼ 60 µm away from the outer wall (**Figure S8**). For an equivalent range of Reynolds numbers, no rigid spherical or non-spherical particles could mimic that behaviour. Even when a suspension of beads and *L. mexicana* parental cells was generated, both the parasites and the beads maintained their respective focusing positions towards the inner wall and outer wall (**Figure S9**). These result might indicate the presence of smaller secondary Dean vortices near the outer wall, arising from the non-spherical cells deforming the fluid profile [17]. The presence of secondary vortices has been demonstrated experimentally via confocal microscopy and shown to trap both rigid and deformable particles at the outer wall [41]. Unfortunately, this approach could not easily be replicated here due to the thickness of the materials used for the microfluidic devices to withstand the flow pressure (layers ∼ 1 mm thick). Acquiring good quality and meaningful images of single cells passing into the channel at such high Reynolds numbers is challenging from an experimental standpoint, which further highlights the needs for accurate modelling tools to mimic such experiments in devices with a constantly changing radius of curvature such as spiral designs. More work is currently underway to better understand this generic focusing towards the outer wall but the approach can nevertheless be translated to a range of applications where *e*.*g*. cell concentration is needed or cell sorting. The work highlights how important an accurate size and shape characterisation is for IM experiments. The field of microfluidics and IM in particular has been significantly enhanced by the field of (imaging) cytometry [42-45] and this should be used, when possible, in tandem with IM characterisation of cells to provide the most accurate representation of focusing patterns. We also further demonstrate that the relationship between particle shape, rotation and equilibrium position within curved flows is complex, highlighting the need for further studies into the rotational behaviours of non-spherical particles in curved channels.

## Authors contribution

JH, NH, SO, TH and MJ designed the experiments, JH, NH and SO performed the experiments. JH designed the data processing approach and analysed the experimental results with MJ. JH and MJ wrote the manuscript; all authors discussed the results and commented on the manuscript.

## Acknowledgements

JH and NH would like to thank the Engineering and Physical Sciences Research Council (EPSRC) for their scholarships (EP/R513222/1 and EP/W524670/1). This project was supported by the Royal Academy of Engineering under the Research Fellowship programme (RF\201718\1741). MJ would like to thank the Royal Society for their support (RGS\R1\191188). SO is funded by a Cunningham Trust (www.cunninghamtrust.org.uk) PhD studentship grant (PhD-CT-19-14) awarded to TH.

