## Supporting figures for "Impact of flagellated and elongated morphological phenotypes on the focusing behaviours of biological cells in inertial microfluidic devices"

### Supplementary information

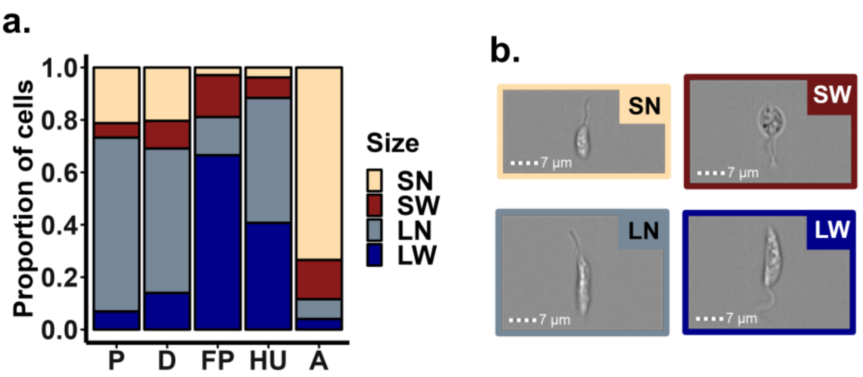


**Figure S1. Distributions of cell shape.**
Two thresholds were applied to provide a deeper outlook into the cell morphologies present in the parental (P), deflagellated (D), flavopiridol- (FP) and hydroxyurea-treated (HU) cells, and amastigotes. These thresholds correspond to the cell length above or below 7.5 µm (which corresponds to one quarter of the channel height), and cell width above or below 4 µm (provides a $\lambda > 0.07$). (a.) For each cell population, every cell was classified into a short (≤ 7.5 µm long) and narrow (< 4 µm wide) (SN), short and wide (≥ 4 µm wide) (SW), long (> 7.5 µm in length) and narrow (LN) or long and wide (LW) category ($n$ > 12,962). (b.) Examples of morphologies in each category, demonstrated with parental cells. Scale bars: 7 µm.

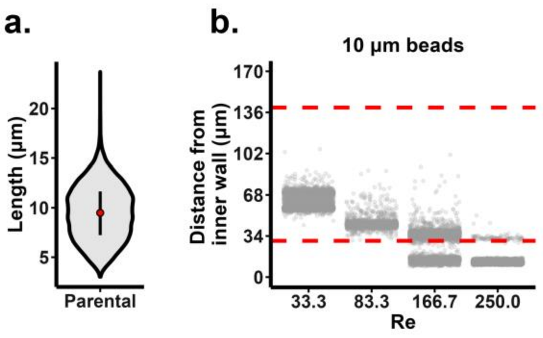


**Figure S2. Beads with an equivalent diameter to *L. mexicana* focus towards the inner wall.**
(a.) The distribution of lengths of fixed *L. mexicana* cells, with a mean of 9.5 µm. The violin plot shows the mean as the red circle and the black bar the 25th - 75th percentiles. (b.) The

focusing position of spherical 10 µm beads within a 30 x 170µm$^2$ channel as analysed using a high-speed camera. Each individual dot corresponds to a single particle imaged in the channel. Red dashed reference lines indicate 30 µm from the inner and outer wall.

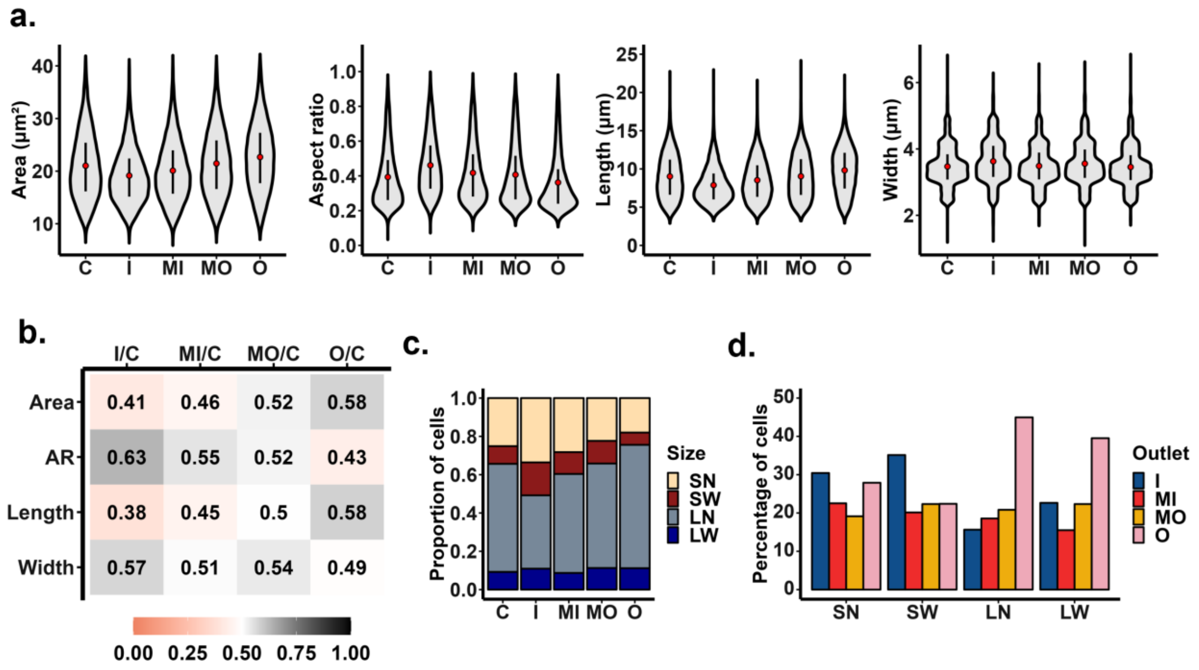


**Figure S3. Sorting of live *L. mexicana* enriches for short and wide cells.**
Live parental *L. mexicana* were sorted at Reynolds number Re = 116.7, and the sorted populations were collected at the four outlets of a spiral IM device with a channel cross section of 30 x 170µm$^2$. Imaging flow cytometry analysis was carried out on an unsorted control (control) and the populations collected from the inner (I), middle inner (MI) middle outer (MO) and outer (O) outlets after sorting. (a.) The area, aspect ratio, length, and width were plotted as a violin plot for each population, with the red circle showing the mean and the black bar the 25$^{th}$ - 75$^{th}$ percentiles ($n \geq$ 18,539). (b.) Comparison of the area, aspect ratio (AR), length and width distributions between the unsorted control population and the each of the different outlets, calculated from the area under the curve.  A score close to 0.5 minimum represents a minimum deviation from the unsorted control population, a value of < 0.5 indicates values smaller than the control while a value > 0.5 demonstrates a value larger than that of the control. (c.) The distribution of sizes of cells sorted into each outlet ($n \geq$ 18,539). The cells from each sample were classified based on their size into short (S, ≤ 7.5 µm in length) or long (L, > 7.5 µm in length) and narrow (N, < 4 µm in width) or wide (W, ≥ 4 µm in width). (d.) The percentage of cells of each size category sorted into each outlet ($n \geq$ 18,539).

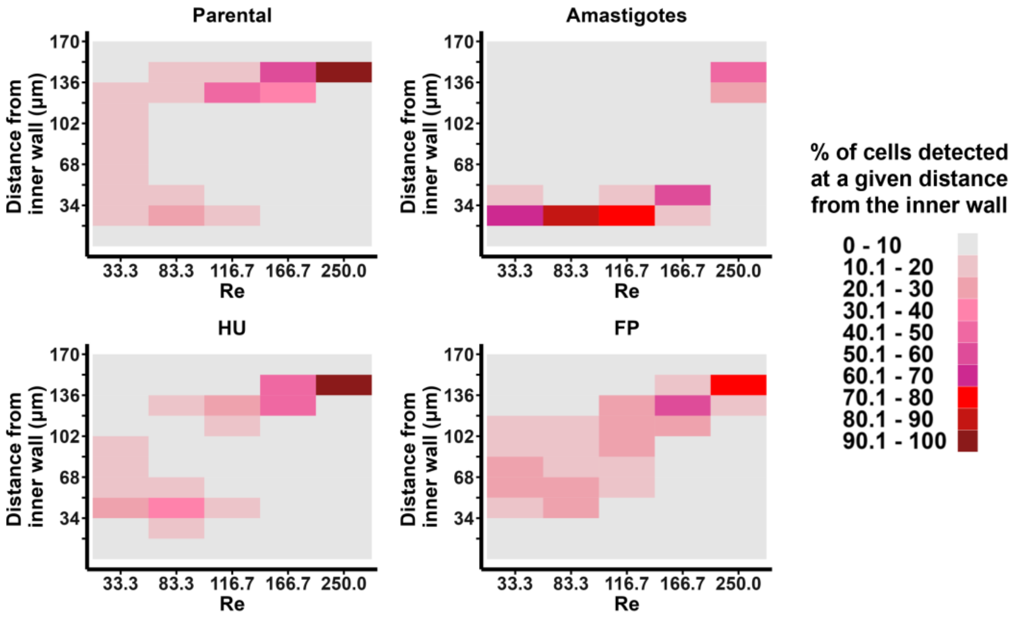


**Figure S4. Characterisation of the focusing behaviour of parental, amastigote, flavopiridol-treated (FP) and hydroxyurea-treated (HU) cells in a spiral inertial device.** High-speed imaging was carried out for each of the parental, amastigote, hydroxyurea- and flavopiridol-treated cell populations, to extract the focusing position of individual particles within the 30 x 170 µm$^2$ channel. From these data, the percentage of cells focusing within 10 regions of 17 µm was calculated for each flow rate and colour coded based on the percentage. Red zones highlight a high focusing concentration with >70% of cells found within a tenth of the channel width. Re denotes the Reynolds number ($n \geq 1{,}000$).

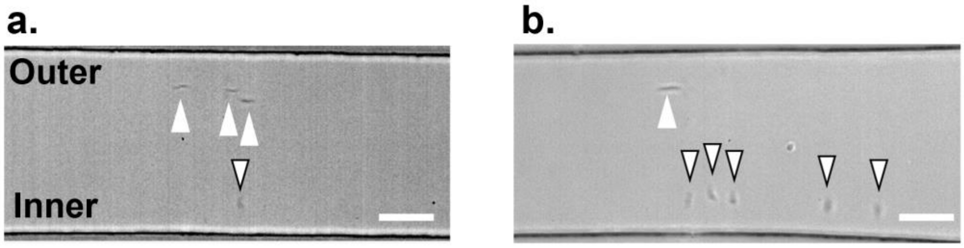


**Figure S5. *L. mexicana* have different orientations at the inner and outer wall.** High-speed images of *L. mexicana* parental cells (a.) and hydroxyurea-treated parasites (b.) focusing at a Reynolds number of 83.3 within the 30 x 170 µm$^2$ channel. Cells which are focused to the outer wall and aligned with the flow are identified with a solid white arrow while cells focusing towards the inner wall and oriented perpendicular to the channel walls are marked with bordered arrows. Scale bars: 50 µm.

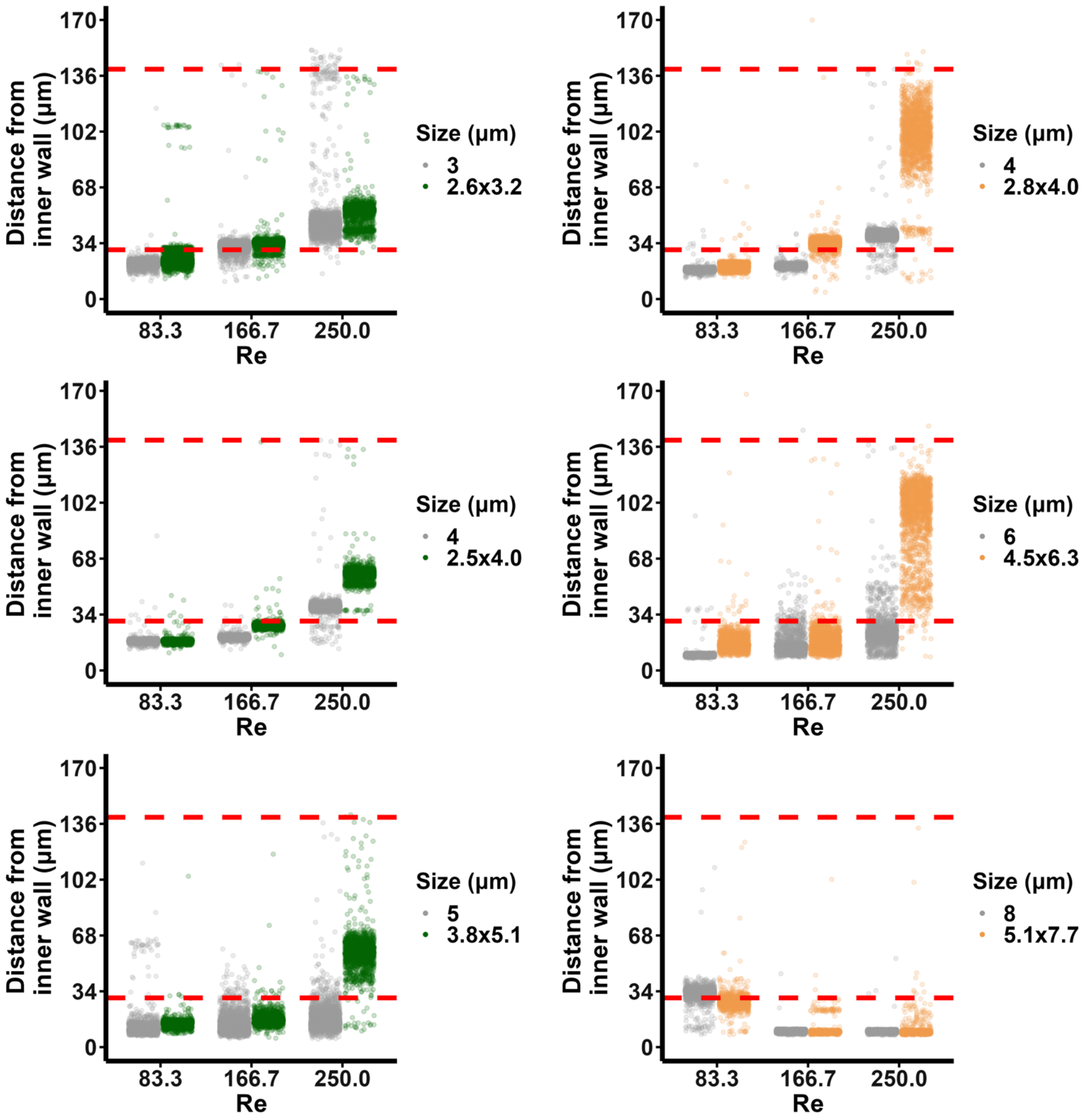


**Figure S6. Characterisation of the focusing behaviour of spherical and non-spherical rigid particles.**

Left graphs: focusing behaviour of pear-shaped beads of sizes 2.6 x 3.2 $\mu m^2$, 2.5 x 4.0 $\mu m^2$ and 3.8 x 5.1 $\mu m^2$ compared to that of spherical 3, 4 and 5 µm diameter beads, respectively. Right graphs: focusing behaviour of peanut-shaped beads of sizes 2.8 x 4.0 $\mu m^2$, 4.5 x 6.4 $\mu m^2$, and 5.1 x 7.7 $\mu m^2$ compared to spherical 4, 6 and 8 µm diameter beads, respectively. All sorting was carried out in the 30 x 170$\mu m^2$ device, showing a single representative replicate for each sample. The spherical beads are shown in grey, the pear-shaped beads in green and the peanut-shaped beads in orange. Red dashed reference lines indicate positions 30 µm away from both the inner and outer walls ($n \geq 1{,}500$).

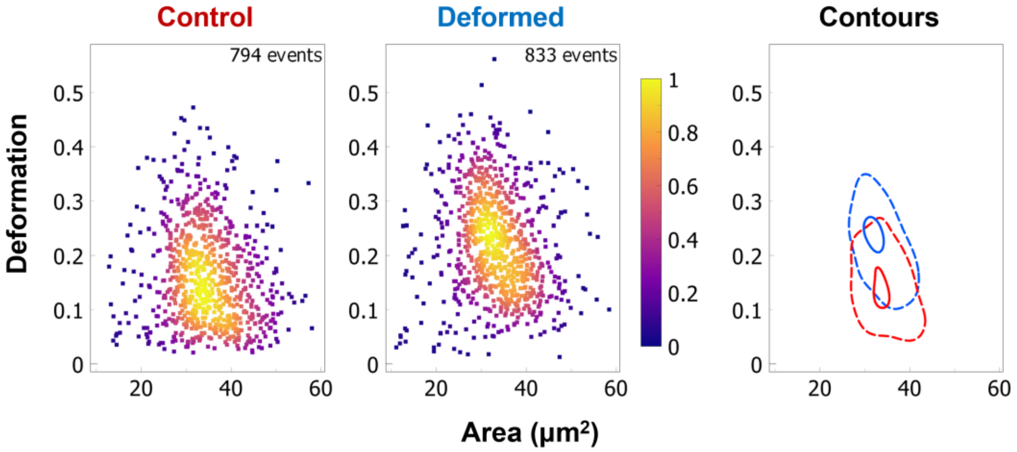


**Figure S7. Flavopiridol-treated parasites deform under shear stress.**
Fixed *L. mexicana* treated with flavopiridol were analysed by deformability cytometry. The density plots display the deformation of individual cells as measured without (control) or with (deformed) shear stress. The densities of the data were calculated using a Gaussian kernel density estimate. The plot on the right-hand side (contours) shows the extracted contours from the density plots, with the deformed contour shown in blue and the control contour shown in red. The dashed and solid lines represent the 95$^{th}$ and 50$^{th}$ percentiles of the data, respectively. A single representative replicate of three is shown (*n* ≥ 794).

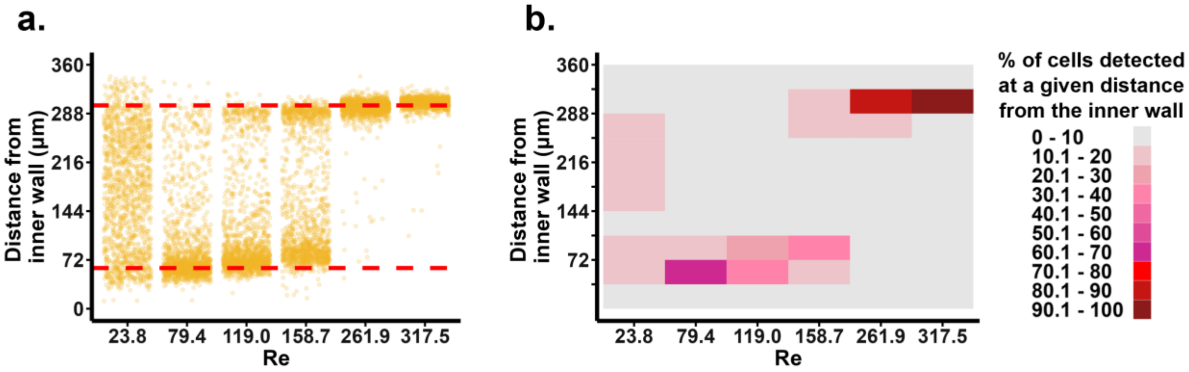


**Figure S8. Fixed parental *L. mexicana* focus to the outer channel in a larger device.**
Fixed *L. mexicana* were analysed for their focusing position in a channel with a cross section 60 x 360 µm$^2$. (a.) The focusing position of individual cells (*n* ≥ 1,500) within the microfluidic channel as analysed by a high-speed camera. The red dashed reference lines indicate a position 60 µm away from both the inner and outer wall (corresponds to a quarter of the channel height). (b.) The data from (a.) was quantified to show the distribution of cells along the channel width. For each flow rate, the channel width was divided into 10 sections; each section corresponds to a distance of 36 µm. Each section was colour-coded (on a scale of 1-10) according to the percentage of cells falling within that section. Each colour bracket accounts for a 10% range.

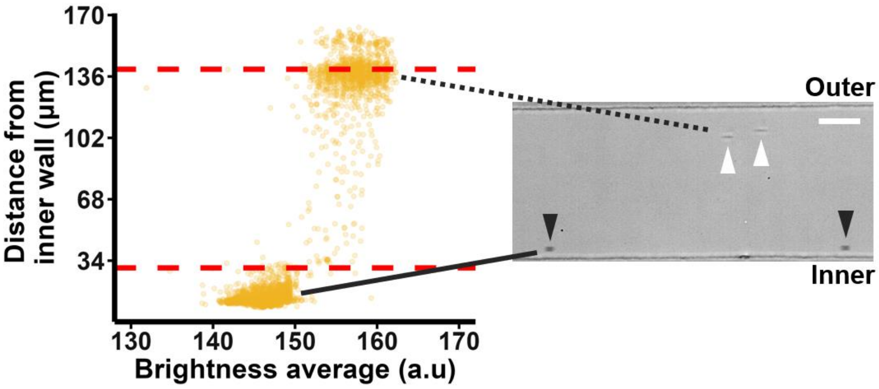


**Figure S9. A mixture of *L. mexicana* parental cells and beads maintain their respective focusing positions.**

A suspension of fixed *L. mexicana* parental cells and beads with a diameter of 5 µm was prepared and the focusing positions of the particles was analysed at a Reynolds number of 166.7 in the 30 x 170 $\mu m^2$ channel. The position of each particle in the channel was plotted against its average brightness (left), and reference lines 30 µm from the inner and outer wall are shown with dashed red lines ($n$ = 5,103). Objects with a brightness ≤ 150 arbritary units (a.u.) were identified as beads, while *Leishmania* cells had a brightness > 150 a.u. An image containing both parental cells (white arrows) and 5 µm beads (black arrows) is given on the right, with the dotted black line identifying the population of parental cells focusing to the outer wall of the channel and the solid black line indicating 5 µm beads focusing to the inner wall. Scale bar: 50 µm.
